# Direct detection of natural selection in Bronze Age Britain

**DOI:** 10.1101/2022.03.14.484330

**Authors:** Iain Mathieson, Jonathan Terhorst

**Author notes:** Correspondence to I.M. or J.T.

## Abstract

We developed a novel method for efficiently estimating time-varying selection coefficients from genome-wide ancient DNA data. In simulations, our method accurately recovers selective trajectories, and is robust to mis-specification of population size. We applied it to a large dataset of ancient and present-day human genomes from Britain, and identified seven loci with genome-wide significant evidence of selection in the past 4500 years. Almost all of them are related to increased vitamin D or calcium levels, and we conclude that lack of vitamin D and consequent low calcium was consistently the most important selective pressure in Britain since the Bronze Age. However, the strength of selection on individual loci varied substantially over time, suggesting that cultural or environmental factors moderated the genetic response to this pressure. Of 28 complex anthropometric and metabolic traits, skin pigmentation was the only one with significant evidence of polygenic selection, further underscoring the importance of phenotypes related to vitamin D. Our approach illustrates the power of ancient DNA to characterize selection in human populations and illuminates the recent evolutionary history of Britain.

## Introduction

Ancient DNA (aDNA) provides direct insight into human evolutionary history. So far, this information has been mainly been used to study demographic history—the migrations, splits and admixtures that our species underwent in the recent past [1]. But in principle, aDNA can also tell us about phenotypic evolution and, in particular, about the contribution of natural selection to phenotypic and genomic variation. Compared to demographic inference, this is more challenging, because studies of natural selection typically require larger sample sizes than studies of population history, which can integrate information from across the genome.

Although some recent studies have used aDNA to study selection and phenotypic evolution, they have mostly focused on a relatively small number of loci [e.g., 2, 3, 4]. Studies that performed genome-wide scans for selection using aDNA [5, 6, 7] have compared allele frequencies across populations, but have not made use of the precise temporal information available from direct dating of ancient samples. For example, the approach in Mathieson et al. (2015) was able to detect selection that happened some time in the past 8,000 years, somewhere in Western Eurasia, but could not be more specific.

With the recent publication of large aDNA datasets [e.g., 8, 9, 7], sample sizes for some regions are now in the hundreds of individuals, large enough to study selection with good spatial and temporal resolution. However, there is a lack of suitable methods to analyze these data. There are many published methods for estimating selection coefficients from time series data [e.g., 10, 11, 12, 13, 14, 15, 16, 17], but all of them unrealistically assume that selective pressures are constant over time, and they are too slow to run on millions of markers at once. We therefore developed a novel statistical approach which is able to estimate arbitrary time-varying selection coefficients, while being fast enough to run genome-wide.

The population of Britain, from 4500 years before present (BP, i.e. the start of the Bronze Age) to the present-day is ideal to demonstrate this approach for several reasons. First, it is relatively homogeneous in terms of both genetics and environment. Second, it is the population with the largest aDNA sample size. Third, there is a large amount of data about the genetic basis of complex traits in this population due to analysis of the UK Biobank. Finally, one of the few studies that attempted to detect selection over this time period (based on data from present-day individuals) was performed in this population [18], giving us a point of comparison for our aDNA-based approach.

## Methods

We begin by formally defining the data, our inference problem, and the model we used to solve it. Our model is a generalization of the one used in [12]. We assume a haploid Wright-Fisher population model whose size *t* generations before present was 2*N_t_*. Time runs backwards, so that *t* = 0 is the present, and *t* = *T* is the earliest point where we have data. We are interested in the frequency of a single allele with two types *A* and *a*. In generation *t*, the *a* allele has relative fitness 1 + *s_t_*/2 relative to *A*. Our inferential target is the vector **s** = (*s*_0_, …, *s_T_*) of selection coefficients over time. The population size history (*N*_0_, …, *N_T_*) is also allowed to vary with time, but we assume that it is known and do not attempt jointly estimate it with **s**.

The data consist of pairs of counts 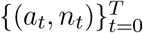. Each pair 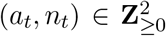 represents the number *a_t_* of *a* alleles observed out of *n_t_* samples collected *t* generations ago. Missing data or generations where no sampling occurred are indicated by setting *n_t_* = 0. Our model conditions on the sample sizes *n_t_*, and we suppress notational dependence on them going forward.

The data generating model is as follows. At time *t*, let the (unobserved) population frequency of the *a* allele be *f_t_* ∈ [0, 1]. Given *f_t_*, the data are binomially distributed with success probability *f_t_*:

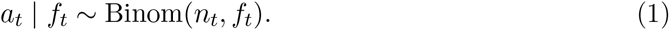

The latent trajectory **f** = (*f*_0_, *f*_1_, …, *f_T_*) evolves according to a Wright-Fisher (WF) model with genic selection and no mutation [e.g., 19]. Given *f_t_* and *s_t_*, the number of individuals *F*_*t*−1_ ∈ [0, 2*N*_*t*−1_] possessing the *a* allele in next generation has distribution

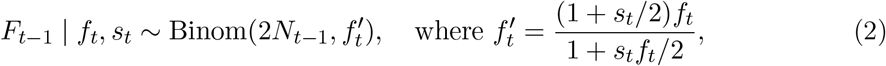

and then we set *f*_*t*−1_ = *F*_*t*−1_/(2*N*_*t*−1_).

Let **a** = (*a*_0_, …, *a_T_*) denote the data. The complete likelihood is

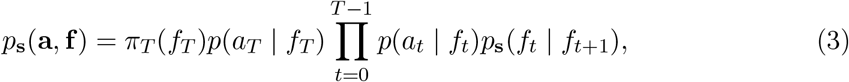

where *π_T_* (*f_T_*) is a prior distribution on the initial allele frequency (described in more detail below), and the probabilities *p*_**s**_(*f_t_* | *f_t_*_+1_) and *p*(*x_t_* | *f_t_*) are specified by (1)–(2). (Throughout this section, we use the notation *p*_**s**_ to denote probability distributions that depend on the selection parameters **s**.) The likelihood of the observed data is obtained by marginalizing (3) over the latent allele trajectory **f** :

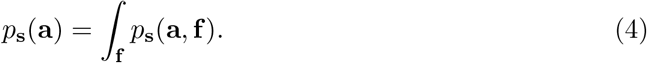

By exploiting the Markov structure of (3), the integral (4) can be efficiently evaluated using the forward algorithm. Each step of the forward algorithm costs 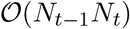 owing the need to evaluate the transition probability (2) for all possible values of *f*_*t*−1_ and *f_t_*. When the effective population size is large (greater than 10^3^, say) this quadratic scaling causes computation to become slow, and it is advantageous to model the latent allele trajectory *f_t_* using a continuous approximation. Several have been proposed, including using the WF diffusion [10, 11, 13, 15, 20]; Gaussian approximations to the WF model [12, 13, 14, 16]; and approximations based on the beta distribution [17, 21, 22]. See also [23] and references therein. A recent review [24] found the beta-with-spikes (hereafter, BwS) approximation of [17] to perform consistently better than other approaches, so we use it as our starting point.

In this model, the latent frequency *f_t_* ∈ [0, 1] of the selected allele is modeled as a mixture distribution with three components. There are two atoms at *f_t_* = 0 and *f_t_* = 1 to allow for the possibility of allele loss or fixation, and the third component is a beta density characterizing the intermediate frequencies, *f_t_* ∈ (0, 1). The form of this model is motivated by the fact that, in the original WF process, the probability of loss or fixation is positive, whereas it is zero if *f_t_* is modeled using an absolutely continuous density.

Although the BwS model is state-of-the-art, room for improvement remains. [24] found that, while generally accurate, the BwS approximation degrades when selection is strong and the effective population size is small. Crucially, we cannot necessarily rule such a regime out when analyzing aDNA data. Another potential shortcoming, not reported by [24] but encountered when we implemented the BwS model, concerns the method used to approximate the transition probability

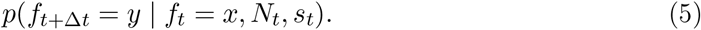

We found that errors in the moment recursions used to compute these probabilities [24, eqn. (6) onwards] tended to accumulate, leading to numerical instability and situations where the variance of the resulting beta approximation, or the spike probabilities, were sometimes computed to be negative. (Python code illustrating this phenomenon is included in the supplement.) This led us to consider refinements of the BwS model.

### Beta mixture model

Breakdown of the moment-based approximation can be explained by insufficient degrees of freedom. The two-parameter beta distribution is not flexible enough to accurately approximate the transition density (5) in all cases. A potential solution is to enrich the approximating class of distributions, by modeling the continuous component of (5) as a *mixture* of beta distributions. This solution is intuitive, and also has theoretical justification: by a famous result of Bernstein [e.g., 25], it holds for any continuous function *g* : [0, 1] → **R** that

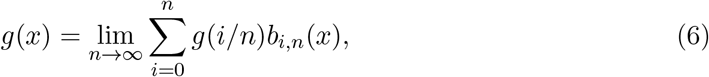

uniformly, where *b_i,n_* is proportional to the Beta(*i* + 1, *n* − *i* + 1) density. Hence, by taking *n* on the right-hand side of (6) to be large but finite, we can accurately approximate any absolutely continuous density on the unit interval. We refer to this model as the *Beta-mixture-with-spikes* (BMwS). A schematic of our model, which accompanies the discussion in the next few subsections, is shown in Figure 1

**Figure 1:**
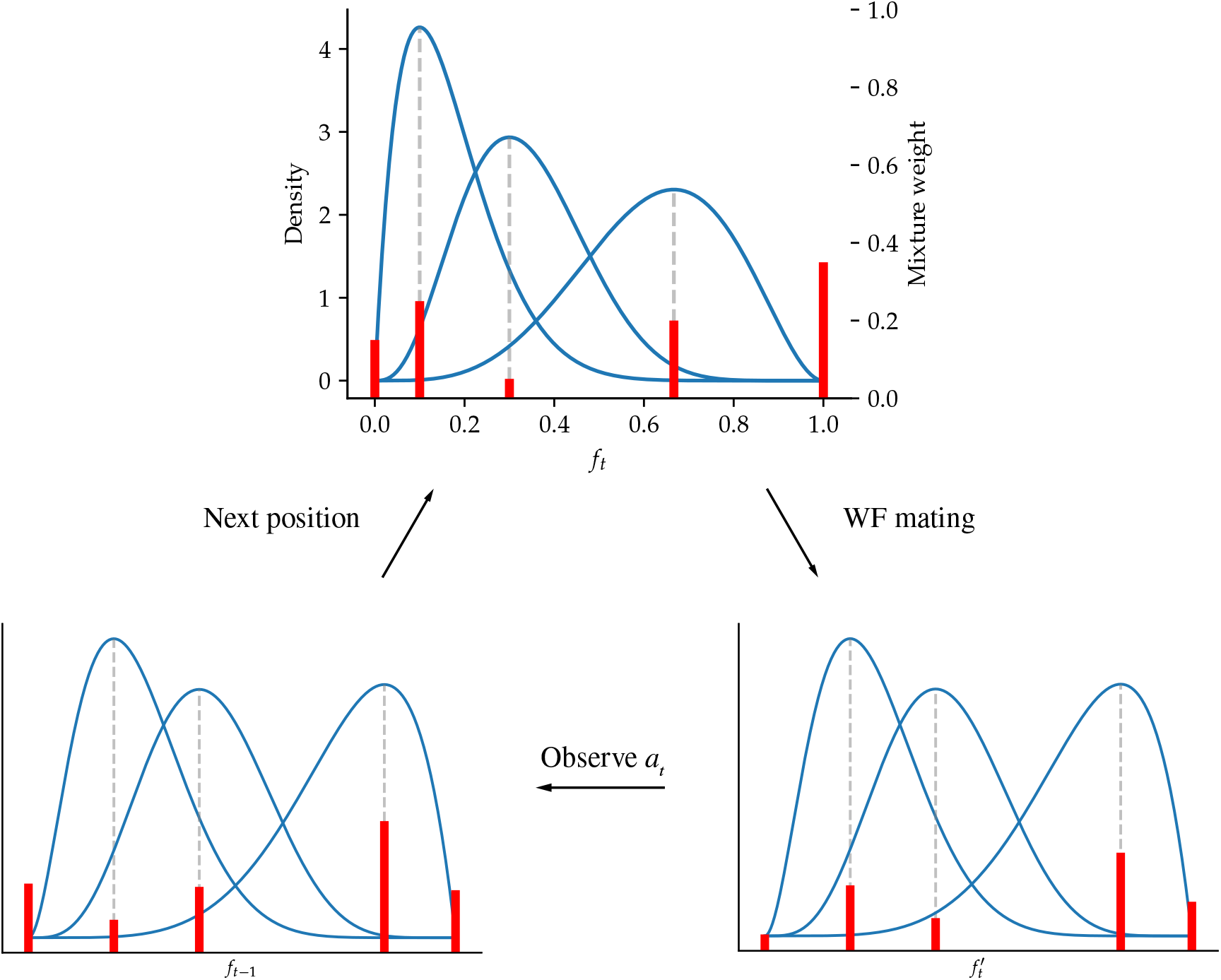
The BMwS model. At each time step *t*, the latent allele frequency *f_t_* is modeled as a mixture of beta distributions, plus spikes at zero and one. In this diagram, there are *M* = 3 mixture components (blue lines). Mixture weights are indicated as red bars, including the spike weights *p*_0_ and *p*_1_ at *f_t_* = 0 and *f_t_* = 1, respectively. After Wright-Fisher mating, the shape of each beta mixture component, as well as the mixture weights, are updated according to equation (13). After observing the data *a_t_*, the mixture weights are again updated according to Bayes’ rule (equation 14). The process then iterates.

Under BMwS, the (posterior) density of *f_t_* is modeled as a mixture of *M* beta densities, plus two atoms at *f_t_* = 0 and *f_t_* = 1. We abuse notation slightly and write this as

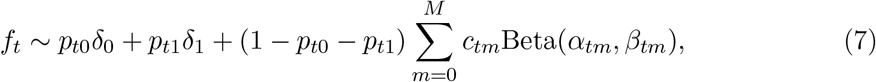

where *δ_x_* denotes a point mass at *x*, and *M* is a user-specified parameter which trades approximation accuracy for speed. To model allele frequency trajectories, we need to characterize the distribution of *f*_*t*−1_ when *f_t_* has the distribution (7). We follow earlier work in utilizing a moment-based approximation, however the form of approximation is new. Previously [e.g., 13, 16, 17, 24], the mean and variance of *f*_*t*−1_ were obtained by Taylor expansion about the infinite-population (zero variance) allele frequency trajectory, and then a moment-matched Gaussian or beta distribution was used to approximate the distribution of *f*_*t*−1_. Here we proceed differently, by directly modeling the action of the WF transition kernel on a beta-distributed random variable.

Assume first that *f_t_* ∼ Beta(*α, β*), and let 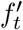 and *f*_*t*−1_ be as in (2). Using a computer algebra system, we determined that

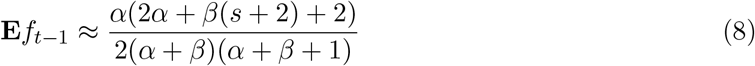

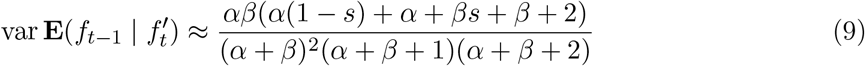

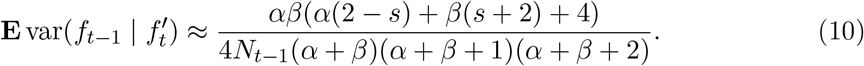

(A Mathematica notebook verifying these computations is included in the supplement.) These approximations are obtained by Taylor expansion of 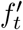 about *s* = 0, followed by substituting in moments of the beta distribution. It would be easy to extend them to higher powers of *s*, but we did not find it necessary, since |*s*| = 0.1 is already at the extreme end of what we expect to find in natural data. Note that for |*s*| < 1 (at least), the above equations imply var *f*_*t*−1_ > 0, so this approximation is robust to the pathology described above.

Using (8)–(10), we can find *α*′, *β*′ such that *f*_*t*−1_ has approximately a Beta(*α*′, *β*′) distribution:

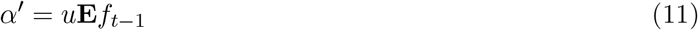

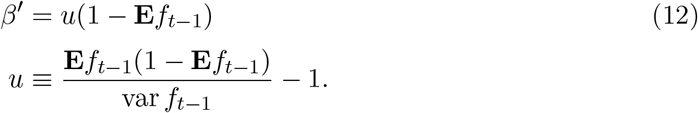

More generally, if the continuous component of *f_t_* is as in (7), then after random mating, we approximate the density of *f*_*t*−1_ by 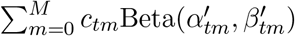.

The other components of the BMwS model are the “spikes” at *f_t_* = 0 and *f_t_* = 1. They are handled similarly to the original BwS model: at each mating event *f_t_* → *f*_*t*−1_, some amount of probability mass is leaked from the beta (mixture) component to atomic components, corresponding to the events *F*_*t*−1_ = 0 and *F*_*t*−1_ = 2*N*_*t*−1_ in (2). (See the next section for a precise statement.) The astute reader will have noticed that, in contrast to some earlier works, we did not properly condition on the complement of this event when constructing the moment approximation shown above. That is, instead of e.g. **E***f*_*t*−1_ in (8), we should instead have considered

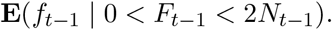

However, the resulting expression is very complex because it involves taking expectation over terms of the form 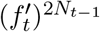. We opted for the simpler and more numerically stable equations (8)–(10), and confirmed in simulations that the model is still accurate across a range of parameter settings.

### Likelihood

The likelihood (4) is calculated using a variant of the usual forward algorithm for hidden Markov models. We explain this computation in greater detail here since the approach is nonstandard.

The forward algorithm recursively updates the so-called *filtering density p*(*f_t_* | *a_t_*, …, *a_T_*), which takes the form shown in equation (7) under our model. Given the filtering density and the observation *x*_*t*−1_, we need to extend the filtering density one step towards the present to obtain *p*(*f*_*t*−1_ | *a*_*t*−1_, …, *a_T_*). This is accomplished in stages:

1. We use equations (8)–(12) to compute 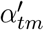 and 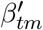, as well as

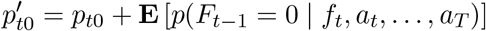

and similarly for 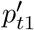. This yields the predictive density

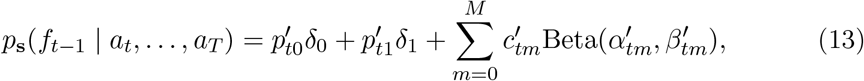

where we defined 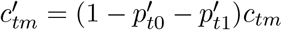.
2. We compute the probability

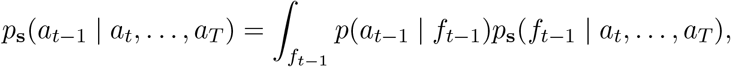

noting that the integral can be evaluated analytically using equations (1) and (13), and conjugacy.
3. We update the mixture weights in (13) to incorporate the information added by observation *a*_*t*−1_. Viewing *a*_*t*−1_ as a draw from the Bayesian hierarchical model defined by (1) and (13), the posterior mixture weights on the beta mixture components are

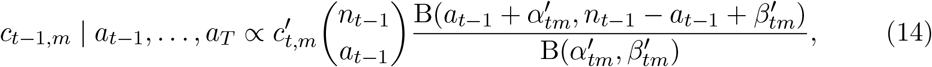

where the right-hand side is the BetaBinomial(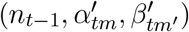 p.m.f. The posterior weight on the atom at *f*_*t*−1_ = 1 is *p*_*t*−1,1_ ∝ **1**_{*a*_*t*−1_=*n*_*t*−1_}_, and similarly for *p*_t−1,0_. The constant of proportionality in these equations is *p*_**s**_(*a*_*t*−1_ | *a_t_*, …, *a_T_*), calculated in step 2.
4. The filtering distribution *p*_**s**_(*f*_*t*−1_ | *a*_*t*−1_, …, *a_T_*) takes the same form as (7), with mixture weights as defined in the preceding step, and 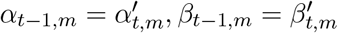.

Recalling equation (4), the log-likelihood of the data is then

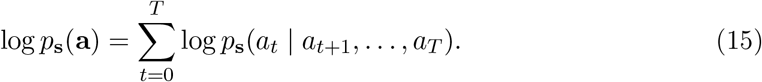

The running time of this algorithm is 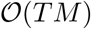, as opposed to the standard forward algorithm which scales quadratically in *M* if it were to denote the number of hidden states in an HMM. This enables us to set *M* fairly large, ensuring that our model can flexibly approximate differently shaped filtering distributions.

### Prior distribution

The filtering recursion is initialized by setting *p*(*f_T_*) = *π_T_* (*f_T_*), where *π_T_* is a prior on the ancestral allele frequency. When developing our method, we observed that the choice of prior affected the accuracy of inferences in the ancient past when analysing aDNA data. This is because when the data are sparsely observed and allele counts are low, there is not enough data to overwhelm the prior in the early stages Markov chain. An uninformative choice of *π_T_* can falsely suggest that the selected allele experienced a large change in frequency, potentially generating a spurious signal of selection. To mitigate this effect, we adopted a coordinate-ascent approach where we alternatively maximized the log-likelihood (15) with respect to a) the selective trajectory **s** and b) the prior distribution *π_T_* . For the prior, we assumed that *π_T_* ∼ Beta(*α_T_, β_T_*) and optimized over *α_T_, β_T_* . Note that, using the interpolation formula (6), we can accurately model an arbitrary prior density if *M* is sufficiently large.

### Inference

Given the probability model and approximations described in the preceding section, inference is now straightforward. Parameter estimation is carried out by as described in the previous section, with one additional modification. Depending on the quality and density of the data, many entries of **s** may only be weakly resolved, and we also found it advantageous to add a regularization term. The objective function for all the analyses reported below was

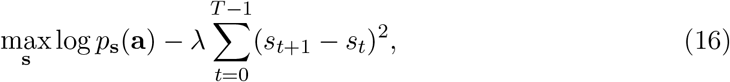

where *λ* > 0 is a tuning parameter. The regularizer penalizes variation in **s**, with larger values of *λ* shrinking all entries towards a single common value, *s*_0_ = · · · = *s_T_* . The number of mixture components for the BMwS model was fixed to *M* = 100, and we performed three rounds of coordinate ascent. For all the examples in this paper, we assumed that the size history (*N*_0_, …, *N_T_*) is known, but co-estimation of selection and size history is also possible using our method, and could be an avenue for future research. Finally, our method is implemented using a JIT-compiled, differentiable programming language [26] to allow for efficient, gradient-based fitting.

### Simulations

We evaluated the performance of the estimator under three different types of selection, each lasting for *T* = 100 generations:

- Constant selection with *s* = 0.01 and initial frequency of 0.1.
- Selection that decreases sinusoidally from *s* = +0.02 to *S* = −0.02 and initial frequency of 0.1.
- Selection that alternates between +0.02 and −0.02 every 20 generations and initial frequency of 0.5.

In each case, we simulated allele frequency trajectories in a Wright-Fisher population with *N* = 10^4^ and then sampled 100 haploid individuals every 10 generations. We also simulated the same selective models under two scenarios of variable effective population size:

- Exponential growth from *N* = 10^4^ to *N* = 10^5^.
- *N* = 10^4^ with a bottleneck of *N* = 10^3^ lasting 10 generations.

For these scenarios we ran the estimator both with the correct effective population size and incorrectly assuming a constant *N* = 10^4^ to evaluate its robustness to misspecification of *N* . We varied the smoothing parameter *λ* from log_10_ (*λ*) = 1 to 6 and report the root mean squared (RMS) bias, variance and total error of the estimator. Finally, we evaluated the error of the estimator as the sample size and frequency vary.

While these simulations cover a plausible range of scenarios for human data (*N* = 10^4^, 100 generations of observations, bottlenecks and exponential growth, selection coefficients of 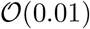), other species may have very different sets of parameters. We therefore recommend that for any particular application, users should run simulations based on their prior parameter values in order to understand the behavior of the estimator in their specific case, and to determine the appropriate smoothing parameter. We provide functions to easily implement simulations under arbitrary selective and population size scenarios. To illustrate this approach, we also ran the simulations described above using the sampling distribution of the British aDNA data used in the rest of paper.

### Ancient DNA data

We collected data from ancient British individuals dated to the past 4500 years from the Allen Ancient DNA resource (v44.3, [27], original sources [28, 29, 30, 31, 7]) and Patterson et al. (2022) [8]. Most samples had been genotypyed on the 1240k array, and the small number of shotgun samples had been genotyped at the 1240k SNPs so we therefore restricted our analysis to this set of SNPs. All data were pseudo-haploid. After removing 22 PCA outliers, we were left with 529 ancient individuals and 98 present-day individuals from the GBR population of the 1000 Genomes project [32], processed into pseudo-haploid data as part of the AADR (Fig. 2A).

**Figure 2:**
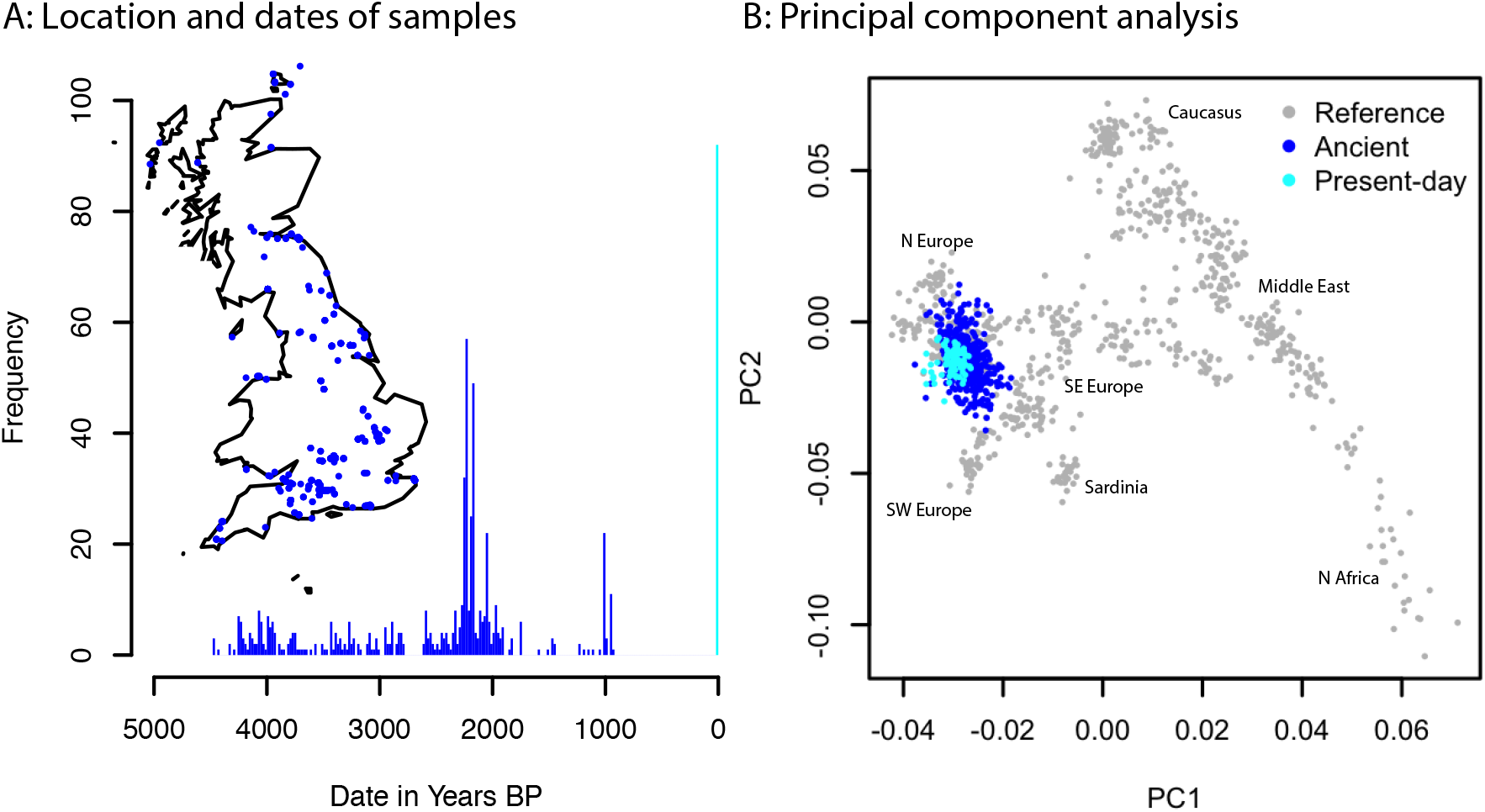
Ancient British data. **A**: histogram of dates of ancient individuals. Inset map shows locations of sites. **B**: Principal components of ancient and present-day samples projected onto axes defined by 777 West Eurasian individuals (see Ref. [33] for details of these individuals).

Our assumes assumes that samples are drawn from a closed, randomly mating population in which every individual experiences the same selective pressures. While no natural population satisfies these conditions, we chose to restrict to Britain dated in the period 4500BP-present because it is the largest aDNA sample from a time and region that comes close to satisfying the assumptions of the model, for the following reasons. First, Britain is a relatively small region (compared to previous Europe-wide studies), meaning that selective pressures are more likely to be shared. Second, we know from previous aDNA studies that the last major change in ancestry in Britain occurred around 4500BP [30]. While recent work has demonstrated more recent Bronze Age migrations into Britain [8], these involve populations that were genetically similar, and inhabited geographically adjacent regions. Third, we confirmed using principal component analysis (PCA) that all the ancient samples in our analysis clustered with the present-day British individuals from the 1000 Genomes project, and more broadly with other Northwestern European individuals in the context of present-day West Eurasia (Fig. 2B).

### Ancient DNA analysis

Starting with 1,150,639 autosomal SNPs, we removed 428,624 with a minor allele frequency (MAF) <0.1 in the full dataset on the grounds that any SNP with a significant frequency shift in this time period would have intermediate MAF. We also removed 101,967 SNPs with greater than 90% missingness and 210,725 SNPs with MAF=0 in the ancient data to leave 409,232 SNPs genome-wide. We inferred selection coefficients 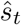 at generation *t* for every SNP in the filtered data using a smoothing parameter of *λ* = 10^4.5^ and an effective population size of *N* = 10^4^. For each SNP, we summarize this estimate by the mean selection coefficient 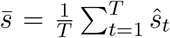, and the root mean squared selection coefficient 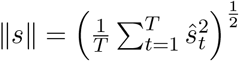. We then computed the mean value of ∥*s*∥ in 20-SNP sliding windows, sliding in 10 SNP increments, so each SNP contributes to two windows. We denote these window statistics ∥*s*∥_20_. Finally, we fit a gamma distribution using the method of moments to the values of ∥*s*∥_20_, and used this fitted distribution to compute P-values for each window. We confirmed that this procedure leads to well-calibrated P-values by repeating the analysis with the dates of each sample randomized, to remove any temporal signal of selection.

We compared our results with two previous genome-wide selection scans. The first—an aDNA based scan [5]—is an allele frequency scan to detect selection in approximately the last 8,000 years in Western Eurasia. The second, the SDS test [18], is a scan based on haplotype lengths in the present-day UK population and is most sensitive to selection in the past few thousand years. First, we restricted the previous scans to the same set of SNPs used in the present scan. Next, for each window in the present scan, we computed the mean test statistics (chi-squared statistic for the aDNA scan and squared Z-score for the SDS) in each window and compared to the test statistics generated by our method.

### Polygenic selection test

We obtained summary statistics from genome-wide association studies (GWAS) for 28 quantitative traits [34, 35, 36, 37, 38]. We took the intersection of GWAS SNPs with the 1240k array, restricted to P-values < 10^−4^, and pruned to an independent set of SNPs by iteratively taking the SNP with the smallest P-value and removing all other SNPs within 250kb. For each trait, we calculated the correlation between effect size and estimated selection coefficient for each independent SNP.

## Results

Our estimator successfully recovers complex selection trajectories from simulated data (Fig. 3), with total sample sizes of the same order of magnitude as our ancient sample, though individual selection coefficient estimates can have considerable uncertainty—on the order of ±0.01 − 0.02 with these parameter values. This suggests that with the data available we should be able to reliably detect selection coefficients around 0.02, similar to the SDS and aDNA allele frequency approaches that we compare with [5, 18]. As expected, the optimal smoothing parameter depends on the true selective trajectory—the smoother the trajectory, the higher the optimal *λ*. We therefore recommend choosing *λ* based on simulations, with parameters that are informed by the specific application.

**Figure 3:**
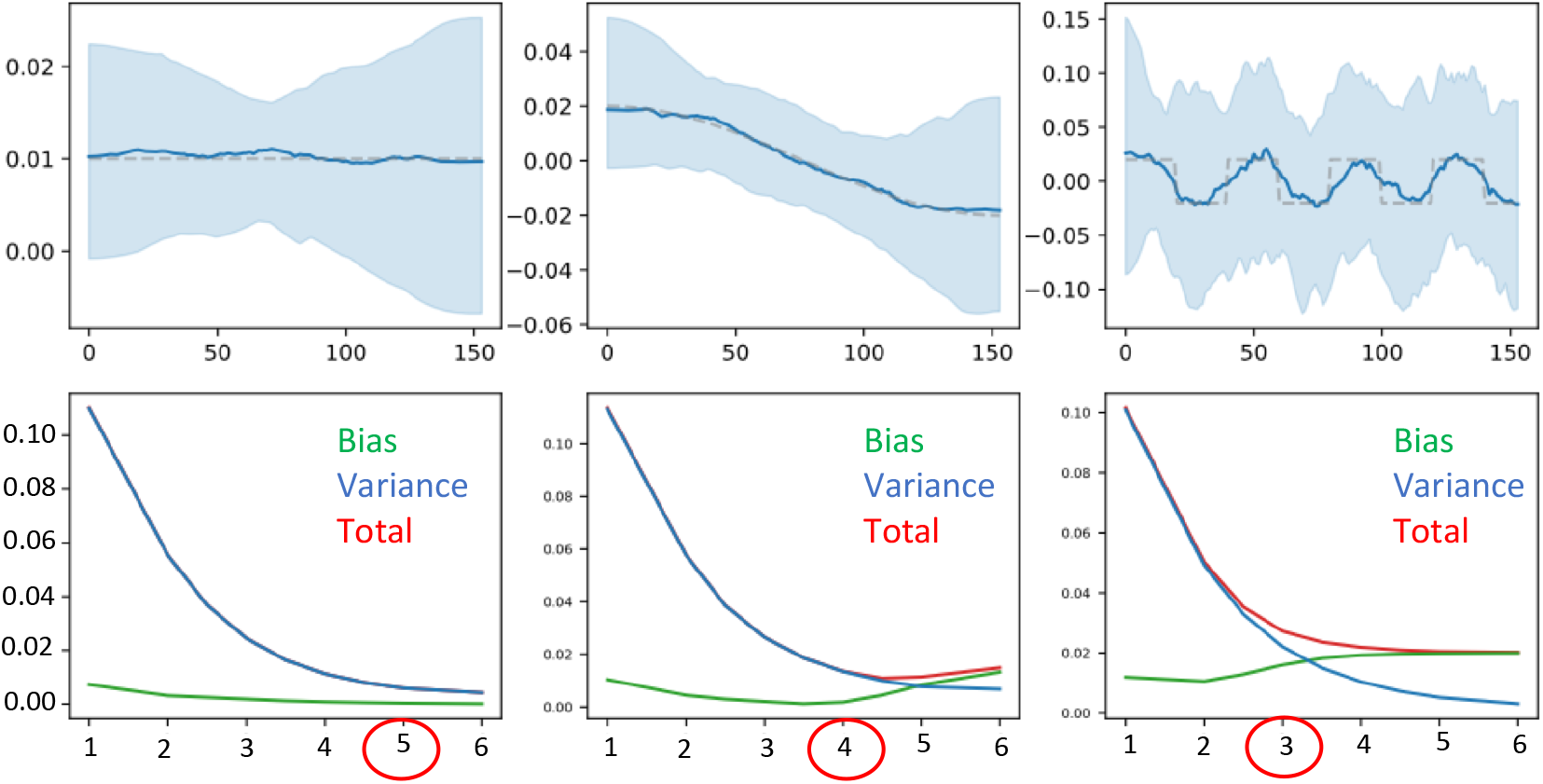
Simulation results. Each column shows a different selection coefficient. **Upper row**: Estimated selection coefficients. Dashed line: simulated selection coefficient. Solid blue line: mean selection coefficient from 100 simulations. Light blue shaded area: region containing point estimates from 95/100 simulations. **Lower row**: Square root of squared bias, variance and total squared error as a function of log_10_ (*λ*). The circled value is the one used for the estimates in the upper row.

The estimator performs similarly in the presence of population size bottlenecks or exponential growth, although it tends to slightly over-smooth changes in *s* in these cases (Fig. S1). Importantly, it is relatively robust to mis-specification of effective population size: if we input constant *N* to the estimator, results are not noticeably different than if we specify the correct population size history (Fig. S2). Increasing size and frequency of sampling reduces error (Fig. S3) though, in general, sample size is more important than sampling frequency and, all else being equal, it is better to have infrequent samples of large sizes than frequent small samples.

In the ancient British data, we identified 7 regions with genome-wide significant evidence of selection (Table 1, Fig. 4, Supplementary Table S2). We used a P-value cutoff of 10^*−*7^ as a genome-wide significant cutoff. While conservative for the 68,061 overlapping windows in our analysis, we used this value because when we reran the analysis with randomized sample dates, no window had a smaller P-value (Fig. 4B). Three of these regions, which we denote HLA1, HLA2, and HLA3 are in the HLA region on chromosome 6 (Fig. S5), though these themselves may contain multiple signals. An eighth apparent signal on chromosome 4 containing the gene *LINC00955* is likely artefactual. The lead SNP rs4690044 has a MAF of 0 in present-day samples but around 0.5 in ancient samples. In gnomAD [39], rs4690044 has a MAF of 0.48, but no homozygotes, suggesting an artefactual call caused by a duplication. An additional locus on chromosome 12 (P=2.5 × 10^−7^) which contains the gene *OAS1* and is known to be a target of adaptive Neanderthal introgression [40] was significant at a Bonferroni corrected significance threshold (P=7.3 × 10^−7^).

**Figure 4:**
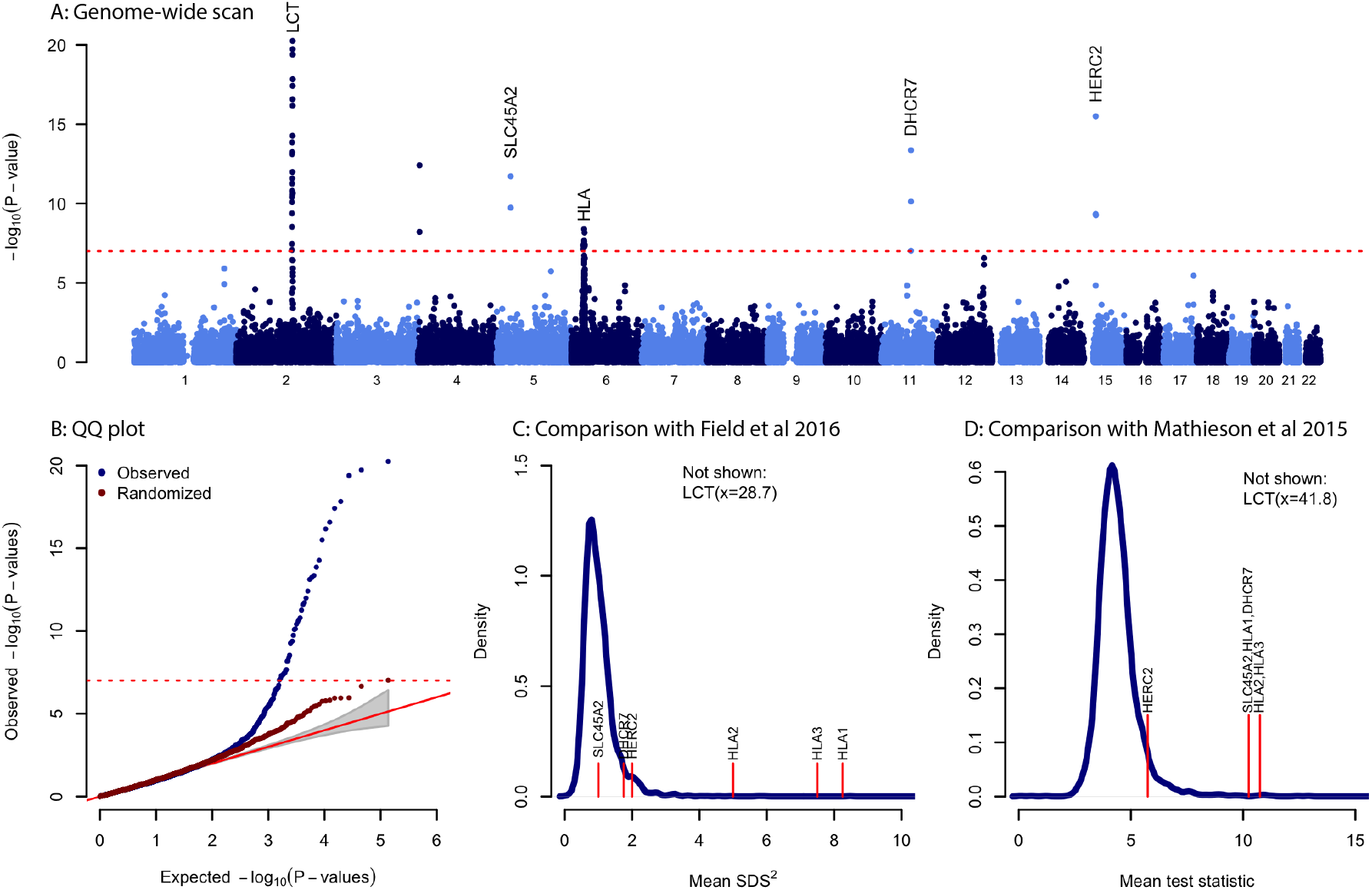
Genome-wide scan for selection in Britain **A**: P-values for selection in 20-SNP sliding windows. Genome-wide significant (P< 10^−7^) windows are labeled with the closest gene or known target of selection. **B**: QQ plot for observations in A (blue) and after randomizing the dates of each sample (red). **C**: Comparison with results of Field et al. [18]. Blue solid line shows the density of mean SDS^2^ in 20-SNP windows. Labeled red lines indicate windows that are genome-wide significant in our analysis. **D**: Comparison with results of Mathieson et al. [5]. Blue solid line shows the density of mean 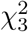 statistic in 20-SNP windows. Labeled red lines indicate windows that are genome-wide significant in our analysis. In both C and D, *HERC2* is approximately at the upper 5^*th*^ percentile of the distribution.

**Table 1:** Genome-wide significant regions. For non-HLA regions, we give the chromosome and position of the center of the lead window, the selected SNP (based on previous literature), the selected allele, and the mean selection coefficient of the lead SNP. For HLA regions we give the co-ordinates of the three regions that contain significant signals (Fig. S5), and the SNPs with the largest test statistic ∥*s*∥. Parentheses indicate that we do not know whether these specific SNPs were the targets of selection.

| Locus | Chr | Start (hg19) | End (hg19) | Target SNP | Allele | $\bar{s}$ |
| --- | --- | --- | --- | --- | --- | --- |
| *LCT* | 2 | 136,582,694 | 136,714,178 | rs4988235 | G | 0.064 |
| *SLC45A2* | 5 | 33,887,419 | 33,964,938 | rs16891982 | C | 0.043 |
| HLA1 | 6 | 28,234,597 | 28,374,902 | (rs6922111) | (C) | (0.046) |
| HLA2 | 6 | 31,026,009 | 31,361,897 | (rs4947296) | (C) | (0.034) |
| HLA3 | 6 | 32,083,175 | 32,581,973 | (rs204994) | (C) | (0.042) |
| *DHCR7* | 11 | 71,142,350 | 71,183,690 | rs7944926 | A | 0.051 |
| *HERC2* | 15 | 28,334,927 | 28,526,228 | rs12913832 | A | 0.017 |

All seven of these regions were identified by a previous allele frequency based selection scan using aDNA to detect selection in West Eurasia over the past 8,000 years [5] (Fig. 4D). Only *LCT* and the HLA region show significant evidence of selection in a haplotype-based scan using present-day sequence data that aimed to detect selection in Britain in the past few thousand years [18] (Fig. 4C) or in a scan based on identifying very recent coalescence times (∼ 50 generations) in the UK Biobank [41].

For the non-HLA signals where we have a strong candidate for the causal variant based on previous literature, we examined the precise timing and trajectory of selection estimates from our model (Fig. 5). The most significant signal was the well-known *LCT* locus on chromosome 2, where the selected allele is associated with lactase persistence [42, 43]. We find that selection for the persistence allele was strongest (*s* ≈ 0.08) from 150 to 100 generations before present (roughly 4500 − 3000BP) before decreasing to around 0.02 in the past 50 generations. This large change in strength of selection might explain the wide range of estimates from models that assume a constant value [3, 43, 44]. At *DHCR7*, the haplotype tagged by the SNP in our analysis, rs7944926, is associated with protection against vitamin D insufficiency [45] and has been shown to have been under recent selection in both Europe [5] and East Asia [46]. We infer that, in Britain, the selection coefficient increased over the past 150 generations, from around 0 to 0.06, leading to a increase in frequency from about 20% to 60% over the past 3000 years (Fig. 5B).

**Figure 5:**
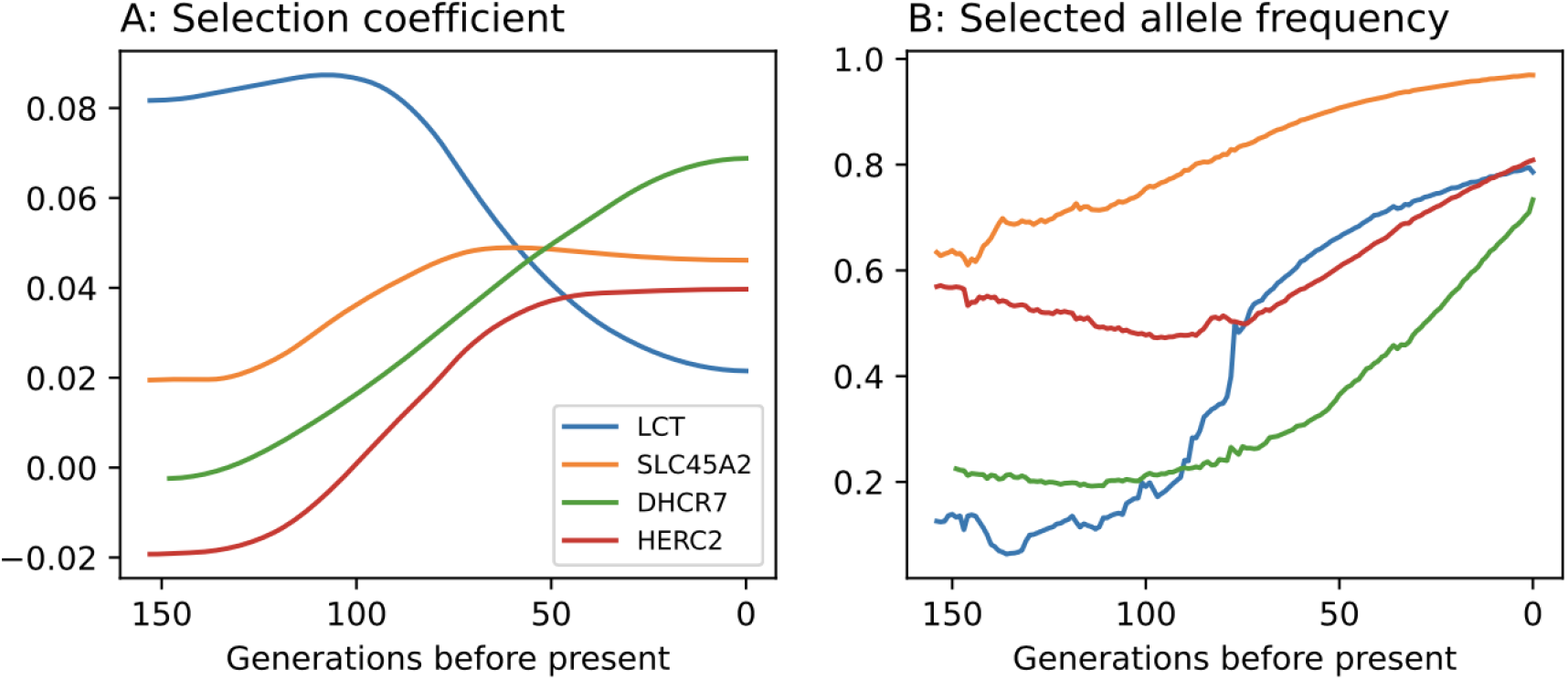
Trajectories of genome-wide significant non-HLA loci. **A**: Estimated selection coefficient. **B**: Estimated allele frequency trajectory.

Derived alleles at *SLC45A2* and *HERC2* are associated with light skin, hair and eye pigmentation [47, 48, 49, 50, 51]. Both these alleles have been shown to have been under selection broadly in Europe [47, 52, 53] and specifically during the Holocene [2, 5]. Time series of aDNA have shown that both alleles were under selection in the past 5000 years in Eastern Europe [2]. There, the derived *SLC45A2* allele increased in frequency from 40% to 90% over the past 5000 years, suggesting a selection coefficient of around 0.03, very similar to our estimate of selection in Britain during the same time (Fig. 5). For the derived *HERC2* allele, we estimate a selection coefficient of 0.02-0.04, similar to that estimated in Eastern Europe, although we find that in Britain selection was largely restricted to approximately the past 2000 years. Wilde et al. [2] also found a derived allele of *TYR* to be under strong selection in Eastern Europe, but we find little evidence that it was under selection in Britain, except possibly before 3000BP (window P-value=0.03, Figure S6).

At the HLA we find three regions with genome-wide significant evidence of selection, which we denote HLA1-3 (Fig. S5). All three correspond to regions identified by Mathieson et al. [5] and also have strong evidence of selection in the Field et al. [18] scan (Fig. 4C). Because of high gene density and complex patterns of linkage disequilibrium in the region, we did not attempt to identify causal genes or variants. However we note that the lead SNP at HLA1 is strongly associated with decreased risk of coeliac disease in UK Biobank [49]. The lead HLA2 SNP is associated with increased risk of ankylosing spondylitis [49] but the region contains the gene *HLA-C*, a variant of which is the strongest known risk factor for psoriasis [54]. Finally, the lead SNP at HLA3 is strongly associated with decreased risk of coeliac disease and psoriasis [49]. These associations suggest that risk of these diseases has been affected by selection, even if they themselves are not the direct targets.

Finally, we searched for evidence of polygenic selection by testing for a correlation between GWAS effect size and selection coefficient for 28 anthropometric and morphological traits (Fig. 6, Table S1). We find significant evidence of polygenic selection for reduced skin pigmentation (P=3.6 × 10^−16^) but none of the other 27 traits (P> 0.04). Though not statistically significant, the largest absolute correlations apart from skin pigmentation is for increased calcium. Notably, we do not detect evidence of selection on any of the phenotypes identified by Field et al. [18] as under selection in Britain in the past 2000 years, including height, infant head circumference and fasting insulin.

**Figure 6:**
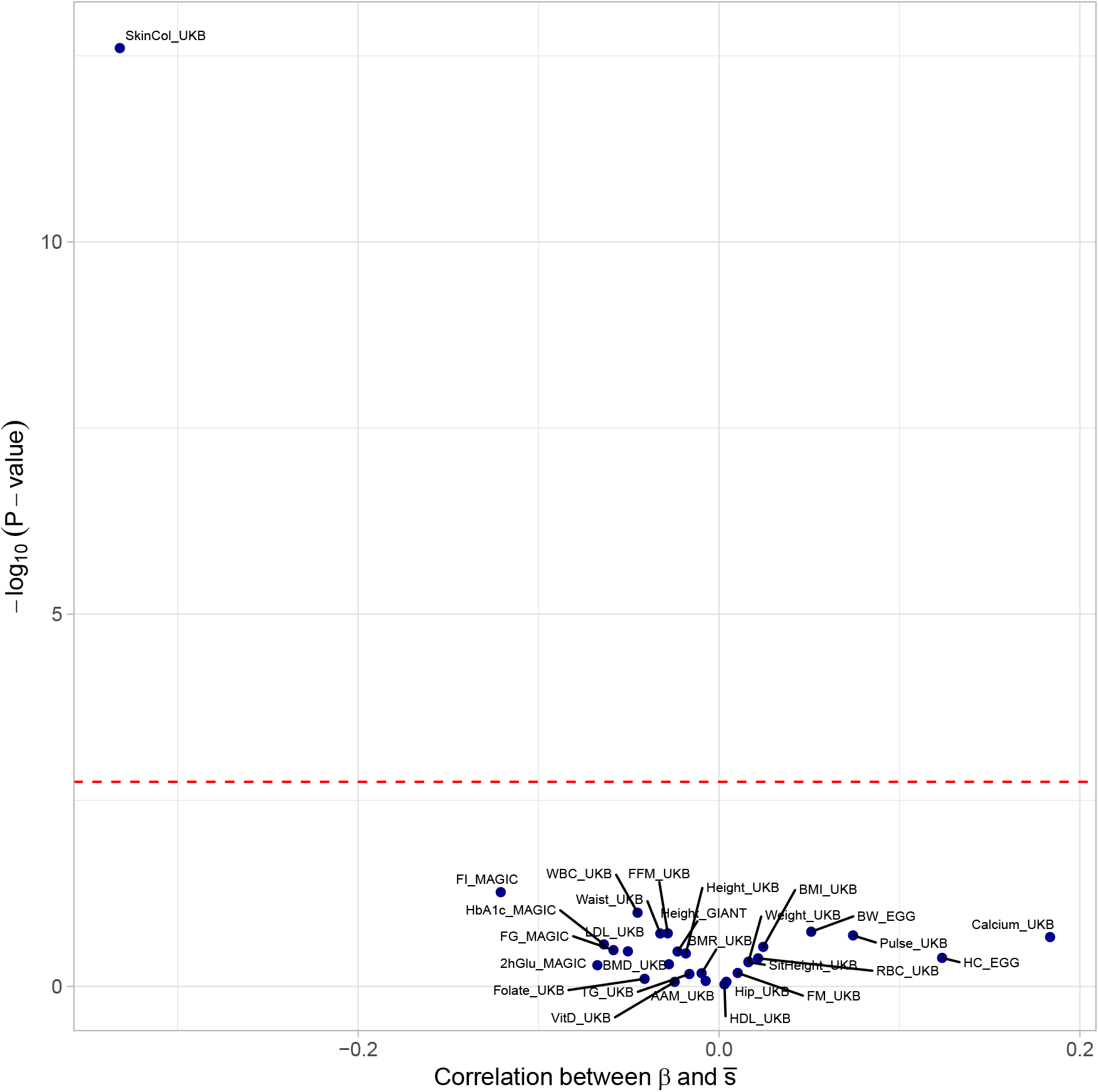
Evidence of polygenic selection. Each point represents a single GWAS. The x-axis gives the (Pearson) correlation between effect size estimates *β* and selection coefficient estimates 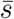 for independent SNPs with GWAS P-value< 10^−4^. The y-axis gives the log^10^(P-value) for null hypothesis of no correlation. Abbreviations, exact values and sources are given in Table S1.

## Discussion

Our fast and flexible estimator allowed us to perform a direct genome-wide scan for selection based on allele frequency trajectories. All of the significant loci we identified have been identified by a previous allele frequency based scan using aDNA [5]. However, that approach scanned for selection broadly in Holocene Europe and was unable to localize selection further in either space or time. Here, we are able to localize selection to Britain in the past 4500 years and, even further, to identify changes in the strength of selection over that time period (Fig. 5). We are also able to identify selected loci that were not identified in studies of much larger samples of present-day individuals [18, 41].

On the other hand, when we search for polygenic selection, the only trait for which we find significant evidence is skin pigmentation, which is known to have been under selection in Europe more broadly into this time period [2, 5, 18, 55]. Though many previous studies [18, 56, 57, 58] reported evidence for recent selection on height, this has been shown to be largely driven by residual stratification in the GIANT consortium GWAS [59, 60]. Evidence of recent polygenic selection for other traits [e.g., 18] likely suffers from the same issue, although UK Biobank GWAS results may be more reliable. Interestingly, even when we test for polygenic selection using the 2014 GIANT height GWAS [36], we find no evidence of selection, suggesting that our approach based on direct estimates of allele frequency changes might be more robust to stratification than others based on haplotype structure or allele frequency differences among extant populations.

While there may have been multiple environmental drivers of selection, taken together our results strongly suggest that the dominant selective pressure in this time was for increased calcium largely moderated through increased vitamin D levels. Vitamin D is required for the absorption of calcium and deficiency leads to bone deformities with potentially major effects on fitness. Since a major source of vitamin D is synthesis in the skin in the presence of UV radiation, the cloudy skies of Britain are likely to have limited this synthesis. The Mesolithic inhabitants of Britain may have avoided this problem through consumption of vitamin D rich marine resources, but later Neolithic and Bronze Age populations including those in our study relied on agricultural products for their subsistence [61], leading to a need for genetic adaptation.

In fact, almost all of our signals of selection can be related to selection for increased vitamin D or directly for increased calcium. Selection at *SLC45A2* and *HERC2*, and selection for lighter skin pigmentation more generally, naturally lead to increased penetration of UV into the skin and therefore higher levels of vitamin D [62, 63]. Lactase persistence allows the consumption of milk which contains both calcium and a small amount of vitamin D. This “calcium absorption” hypothesis has long been suggested to explain the high frequency of the persistence phenotype in Northern Europe [64, 65]. *DHCR7* is directly involved in vitamin D metabolism and the selected allele protects against insufficiency [45]. While the HLA associations are more difficult to specifically identify, two of the selected alleles are protective against coeliac disease which itself is a risk factor for malabsorption of calcium and vitamin D, and consequent osteoperosis [66]. Strong selective pressure for increased vitamin D and calcium levels is therefore a plausible and parsimonious explanation for the patterns of selection that we observe in the this dataset, and suggests that it was the dominant selective pressure in this population.

The main limitation of our approach is the assumption of population continuity. Although there is evidence of external migration into Britain during the time period we investigated [8], there is relatively little change in genetic ancestry since the sources of that migration are genetically similar populations from other nearby parts of Northern Europe. If this affects our results it would likely mean that some of the selection we detected actually occurred in those neighboring populations, somewhat earlier than the dates we find here. However, given that the selection pressures we detect are likely to be shared with these similar populations anyway, we do not think this possibility has a major impact on the interpretation of our results. This limitation is more serious if we want to extend the temporal and geographic range of our analysis. In particular, both ancient and present-day individuals from the Iberian and Italian peninsulas are much more genetically diverse than those from Britain (Fig. S8). Because of this, and the much smaller ancient sample sizes, we did not analyze selection in these Southern European populations, although identifying differences in selective pressures between Northern and Southern Europe is an important direction for future analysis.

Although we have identified the loci under the strongest selection in recent British history, there is evidence in our data of additional selected loci. For example, several known loci including *OAS1* (P=2.5 × 10^−7^, [40]) and *FADS1* (P=1.5 × 10^−5^, [5]) have evidence of selection though below genome-wide significance. With larger sample sizes we can expect that these and other, potentially novel, loci would be identified. Another limitation is that we rely on genotypes from the 1240k array and therefore cannot detect selection on rare SNPs or structural variants that are not tagged by SNPs on the array. Large datasets of shotgun sequence data might extend the range of variation on which selection can be detected.

Our study demonstrates the power of ancient DNA to robustly detect and precisely characterize the timing of natural selection in humans. Notably, with sample sizes of a few hundred, we are able to identify variants under selection that were not identified in studies of thousands of present-day individuals. All the loci that we identify have evidence of substantial change in the strength of selection over the past 4500 years, suggesting that modeling this variability is an important addition to previous approaches that assume constant selective pressure. This also allows us to formulate more specific hypotheses about the environmental drivers of selection. For example, the decrease in the selective advantage of the lactase persistence allele coincides with the Roman occupation of Britain. Perhaps, since cattle milk was not commonly drunk in Roman culture, a culturally mediated decrease in milk consumption in Roman Britain meant that lactase persistence became less useful for obtaining vitamin D, leading to an increase in selective pressures related to other sources (i.e. pigmentation and metabolism). In this way, culture could have determined the genetic response to a constant environmental selective pressure. The prospect of exploring this and similar interactions is an exciting future direction for ancient DNA research.

## Supporting information

Supplemental Notebooks

## Acknowledgments

This research was funded by the NIGMS ([R35GM133708] to I.M.) and the NSF ([DMS-2052653] to J.T.). The content is solely the responsibility of the authors and does not necessarily represent the official views of the funding agencies.

## Data and code availability

Ancient genotype data was from the Allen Ancient DNA Resource v4.3 [27] (original sources cited in main text), and Patterson et al. 2022 [8]. Present-day data from the 1000 Genomes Project [32] was reprocessed as described in the AADR. Code to run the estimator and reproduce the analyses in this paper is available at https://github.com/jthlab/bmws.

---

**Table S1:** Trait descriptions, sources, and values for Figure 6

| Short name | Trait | Ref. | $\rho$ | P | N SNPs |
| --- | --- | --- | --- | --- | --- |
| SkinCol_UKB | Skin Color | [34] | -0.33 | $2.5 \times 10^{-13}$ | 463 |
| Height_UKB | Height | [34] | -0.02 | 0.36 | 2479 |
| SitHeight_UKB | Sitting height | [34] | 0.02 | 0.47 | 1953 |
| Height_GIANT | Height | [36] | -0.02 | 0.34 | 1735 |
| BMI_UKB | Body mass index | [34] | 0.02 | 0.29 | 1836 |
| BMD_UKB | Heel bone mineral density | [34] | -0.03 | 0.50 | 583 |
| BMR_UKB | Basal metabolic rate | [34] | -0.01 | 0.66 | 2028 |
| Weight_UKB | Weight | [34] | 0.02 | 0.46 | 2002 |
| FM_UKB | Fat mass | [34] | 0.01 | 0.66 | 1818 |
| FFM_UKB | Fat free mass | [34] | -0.03 | 0.19 | 2098 |
| Waist_UKB | Waist circumference | [34] | -0.03 | 0.19 | 1592 |
| Hip_UKB | Hip circumference | [34] | 0.00 | 0.87 | 1688 |
| WBC_UKB | White blood cell count | [34] | -0.05 | 0.10 | 1311 |
| RBC_UKB | Red blood cell count | [34] | 0.02 | 0.42 | 1392 |
| Pulse_UKB | Pulse rate | [34] | 0.07 | 0.21 | 291 |
| HDL_UKB | HDL cholesterol | [34] | 0.00 | 0.94 | 656 |
| LDL_UKB | LDL cholesterol | [34] | -0.05 | 0.34 | 363 |
| TG_UKB | Triglycerides | [34] | -0.02 | 0.68 | 631 |
| AAM_UKB | Age at menarche | [34] | -0.01 | 0.84 | 721 |
| BW_EGG | Birthweight | [37] | 0.05 | 0.18 | 677 |
| HC_EGG | Infant head circumference | [35] | 0.12 | 0.41 | 46 |
| VitD_UKB | Vitamin D | [34] | -0.02 | 0.87 | 48 |
| Calcium_UKB | Calcium | [34] | 0.18 | 0.22 | 47 |
| Folate_UKB | Folate | [34] | -0.04 | 0.79 | 44 |
| FI_MAGIC | Fasting insulin | [38] | -0.12 | 0.05 | 254 |
| FG_MAGIC | Fasting glucose | [38] | -0.06 | 0.33 | 285 |
| 2hGlu_MAGIC | Two hour glucose | [38] | -0.07 | 0.52 | 94 |
| HbA1c_MAGIC | Glycated haemoglobin | [38] | -0.06 | 0.27 | 299 |

**Figure S1:**
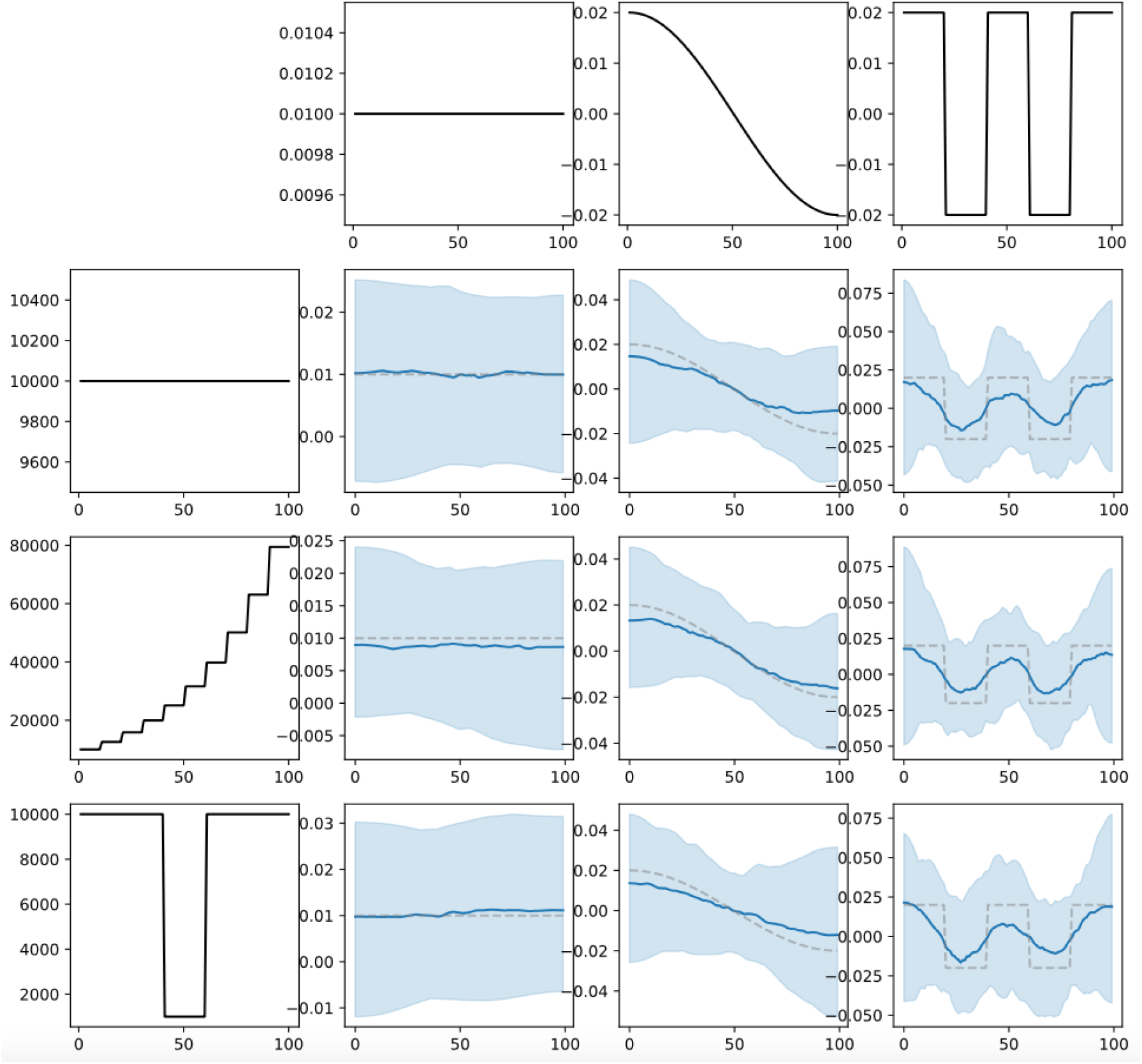
Simulation results for different population size history. Each column represents a different selective trajectory and each row represents a different population size trajectory. Dashed lines: simulated selection coefficient. Solid blue lines: mean selection coefficient from 100 simulations. Light blue shaded areas: region containing point estimates from 95/100 simulations.

**Figure S2:**
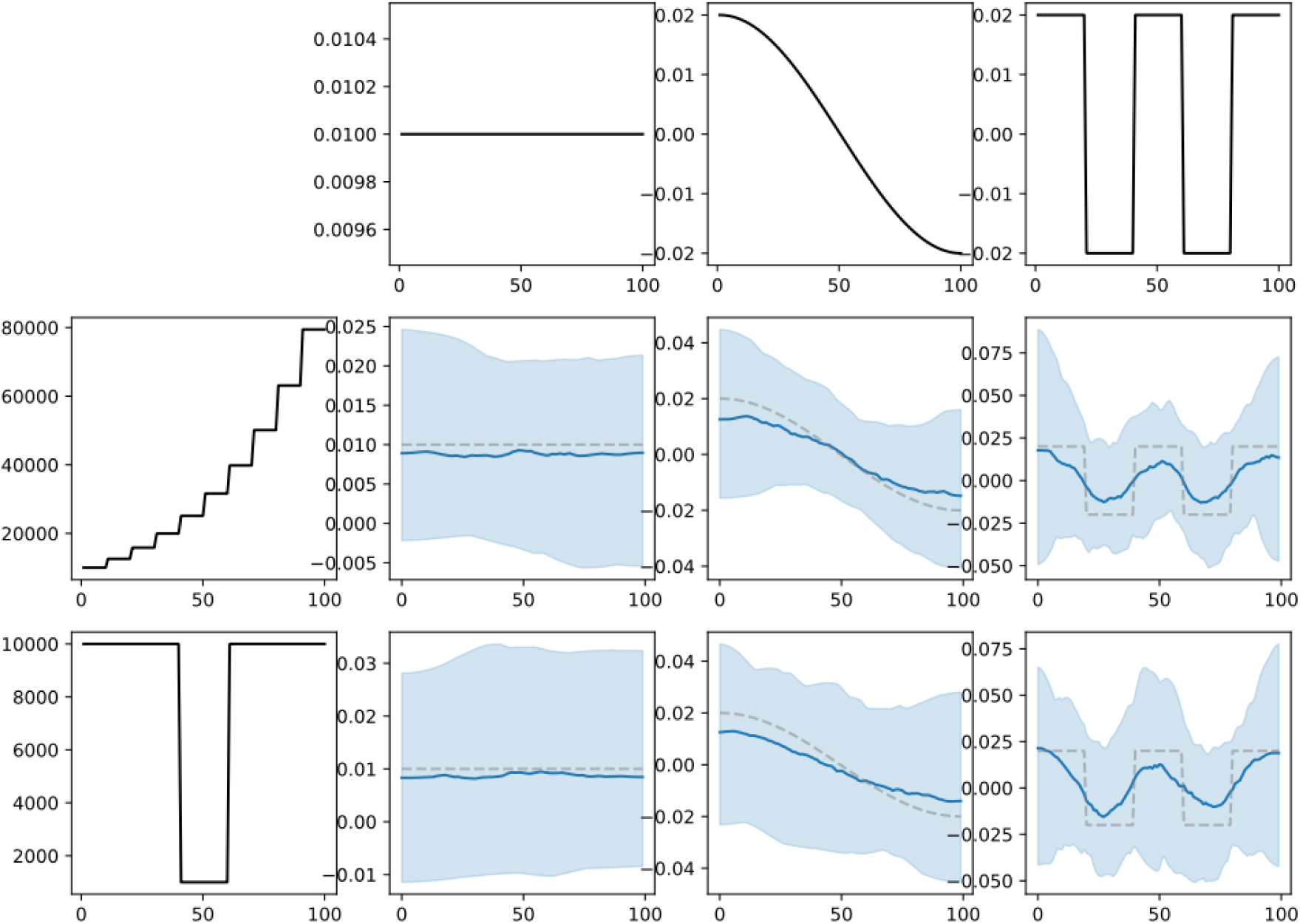
Simulation results for different population size histories. As Fig. S1 but using a constant *N* = 10^4^ in the model, instead of the correct effective population size. Dashed lines: simulated selection coefficient. Solid blue lines: mean selection coefficient from 100 simulations. Light blue shaded areas: region containing point estimates from 95/100 simulations.

**Figure S3:**
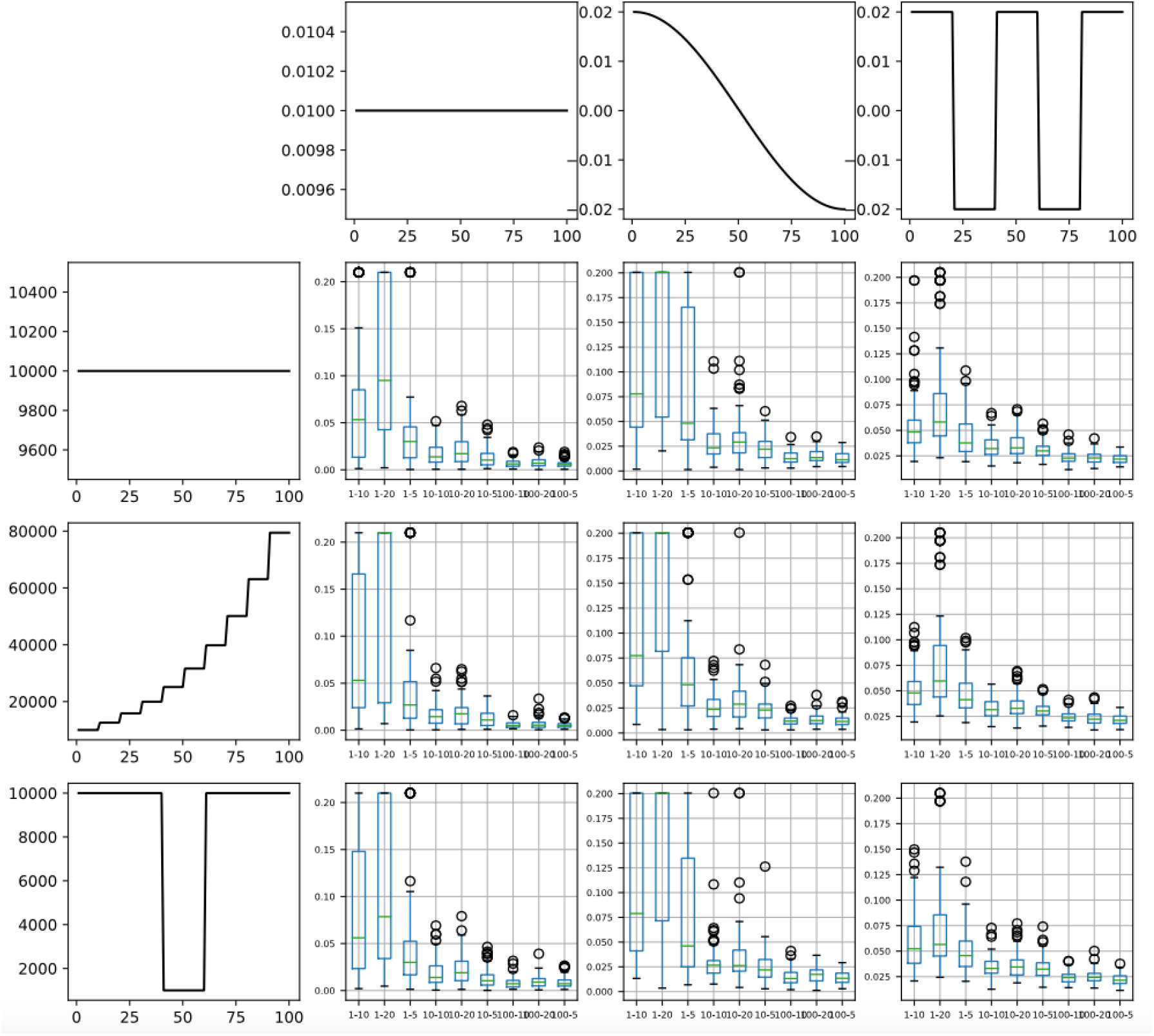
Root mean squared error (RMSE) for different sampling schemes. Each column represents a different selective trajectory and each row represents a different population size trajectory as in Fig. S1. Each barplot shows the distribution of RMSE for 100 simulations based on a different sampling schemes. Sampling schemes are labeled "size-frequency", so for example "1-10" means that we sampled one haploid individual every 10 generations.

**Figure S4:**
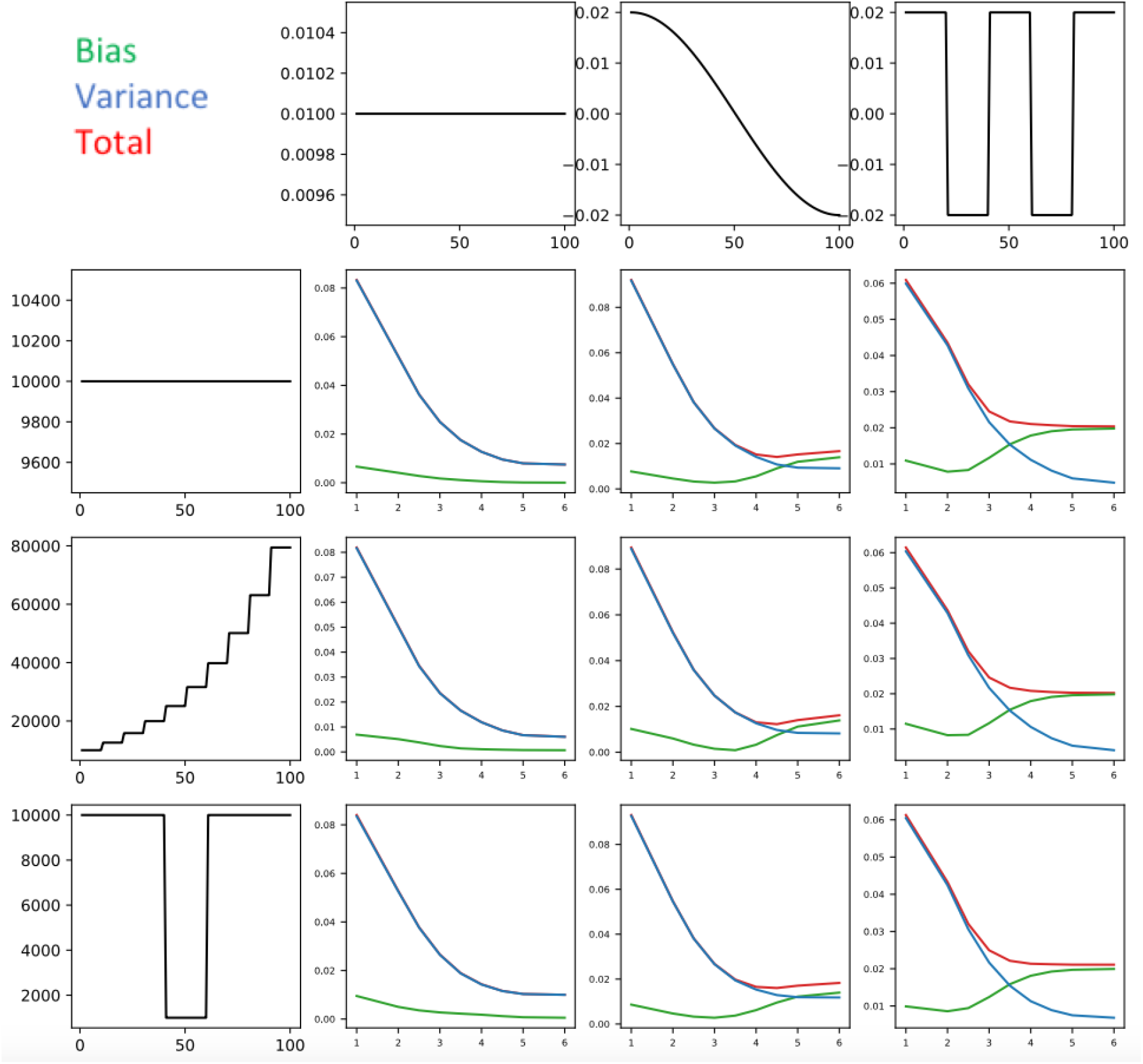
Root mean squared bias, variance and total error as a function of log_1_ 0 (*λ*) using the sampling distribution of the ancient British data for different selective (columns) and effective population size (rows) trajectories.

**Figure S5:**
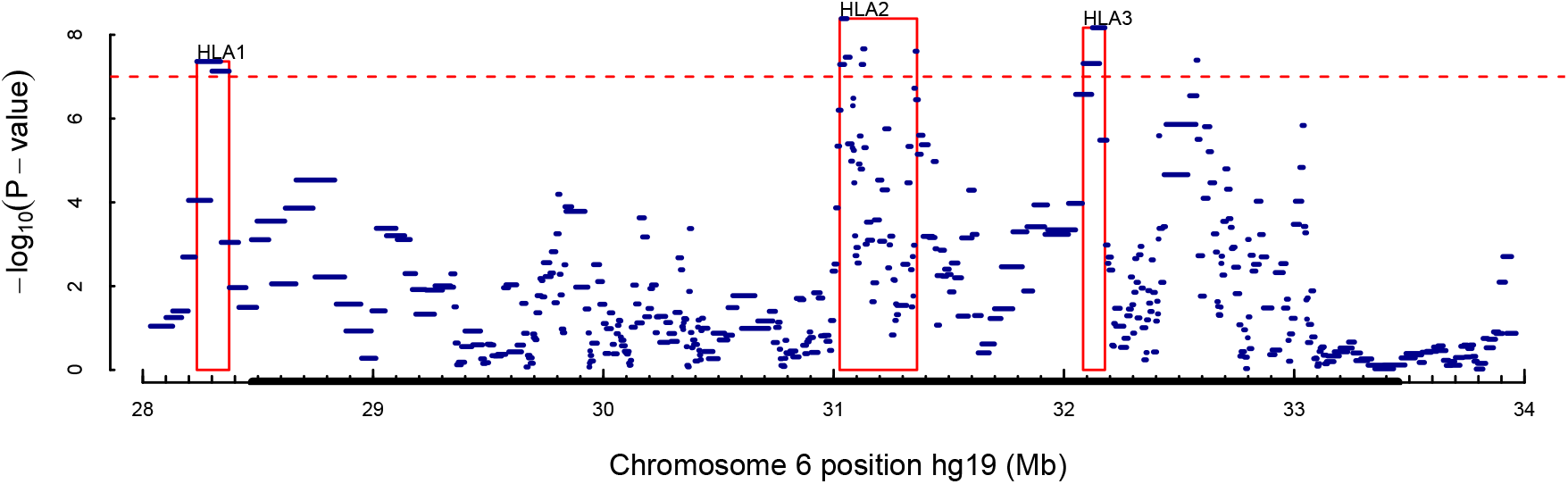
Window-based P-values for the HLA region on Chromosome 6. The thick black x-axis shows the boundary of the HLA. Each blue bar represents a 20-SNP window. We highlight three regions with evidence of one or more significant signals.

**Figure S6:**
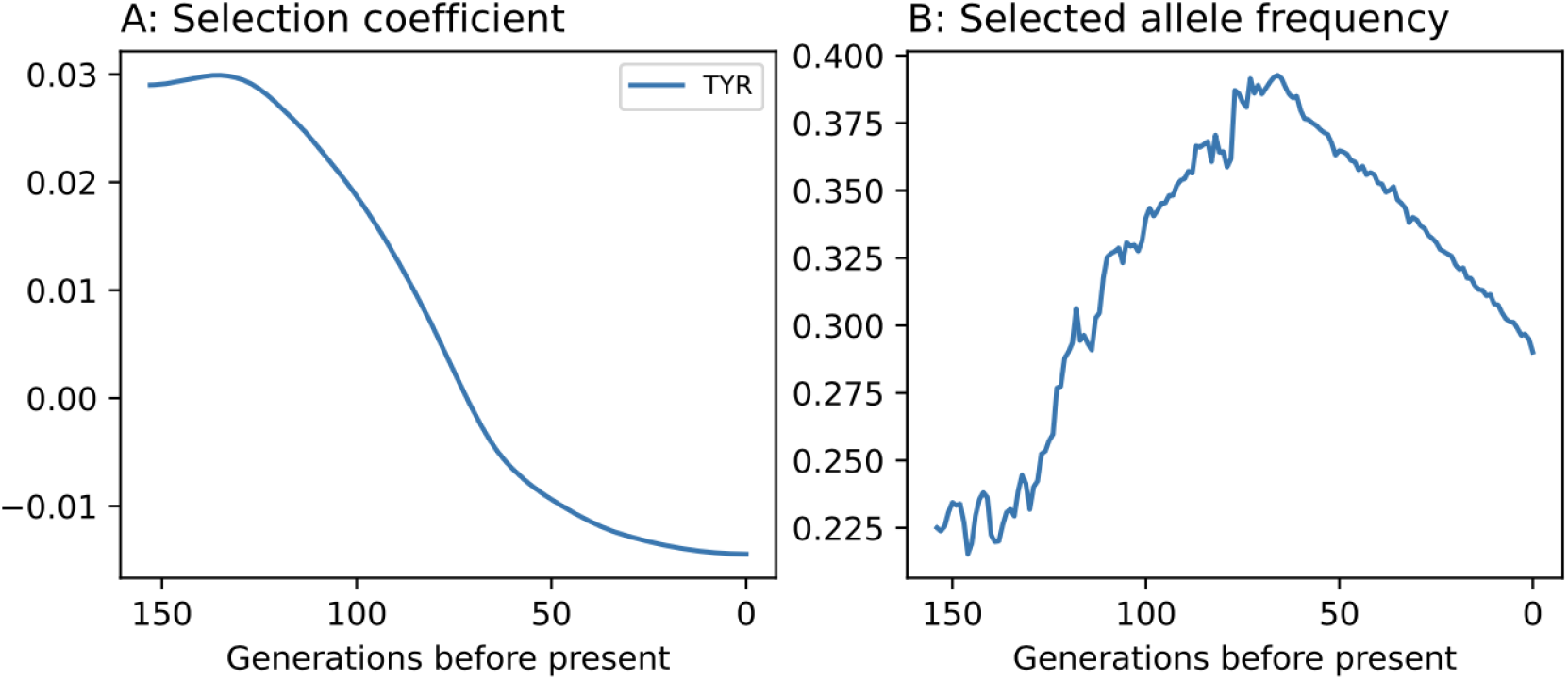
Estimated selection coefficient and frequency for the A allele of rs1042602 at the *TYR* locus.

**Figure S7:**
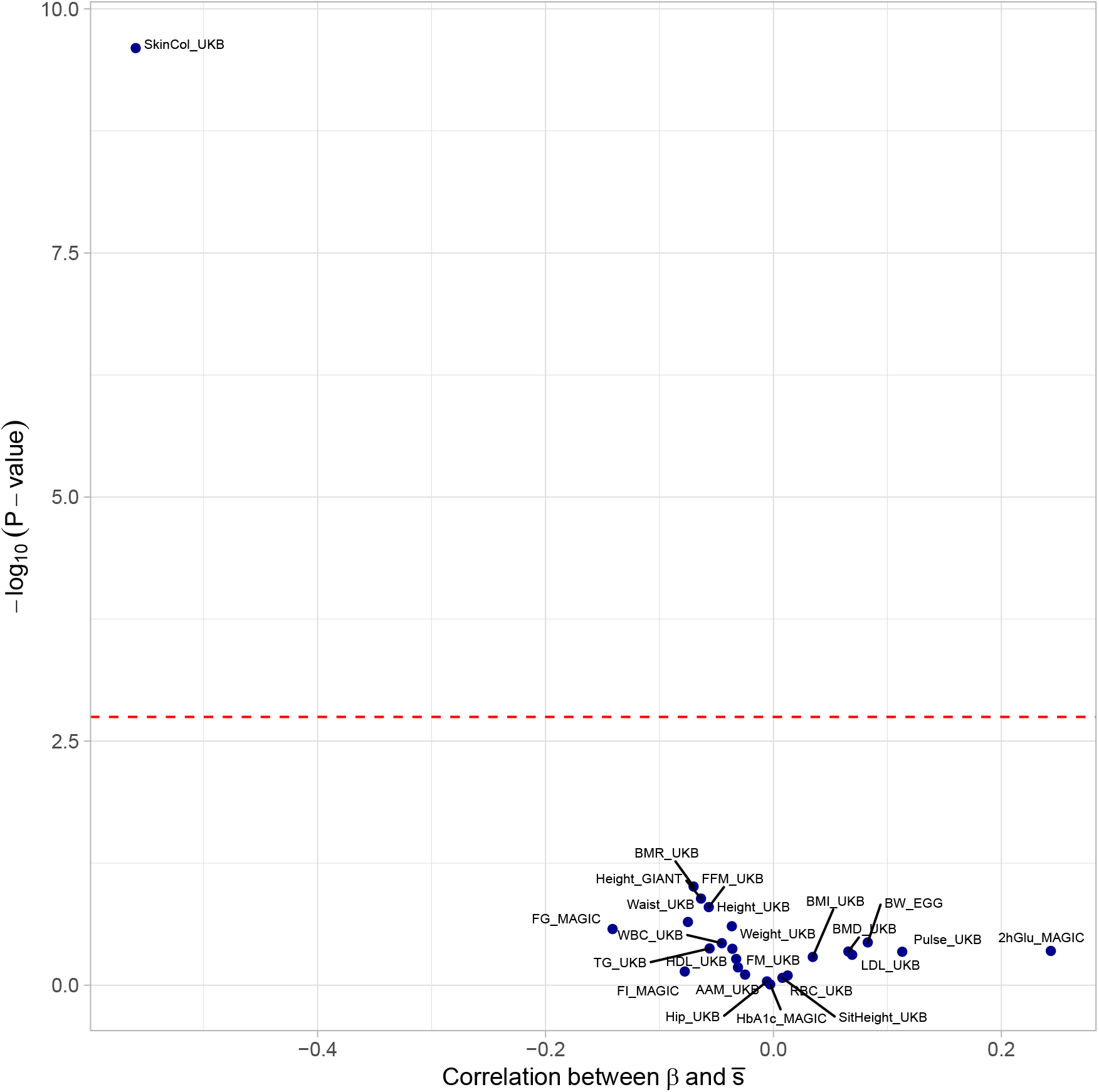
As Fig. 6 but restricting to SNPs that with P< 10^−8^, rather than *P <* 10^−4^. Calcium does not appear on this plot because it has no SNPs with P < 10^−8^.

**Figure S8:**
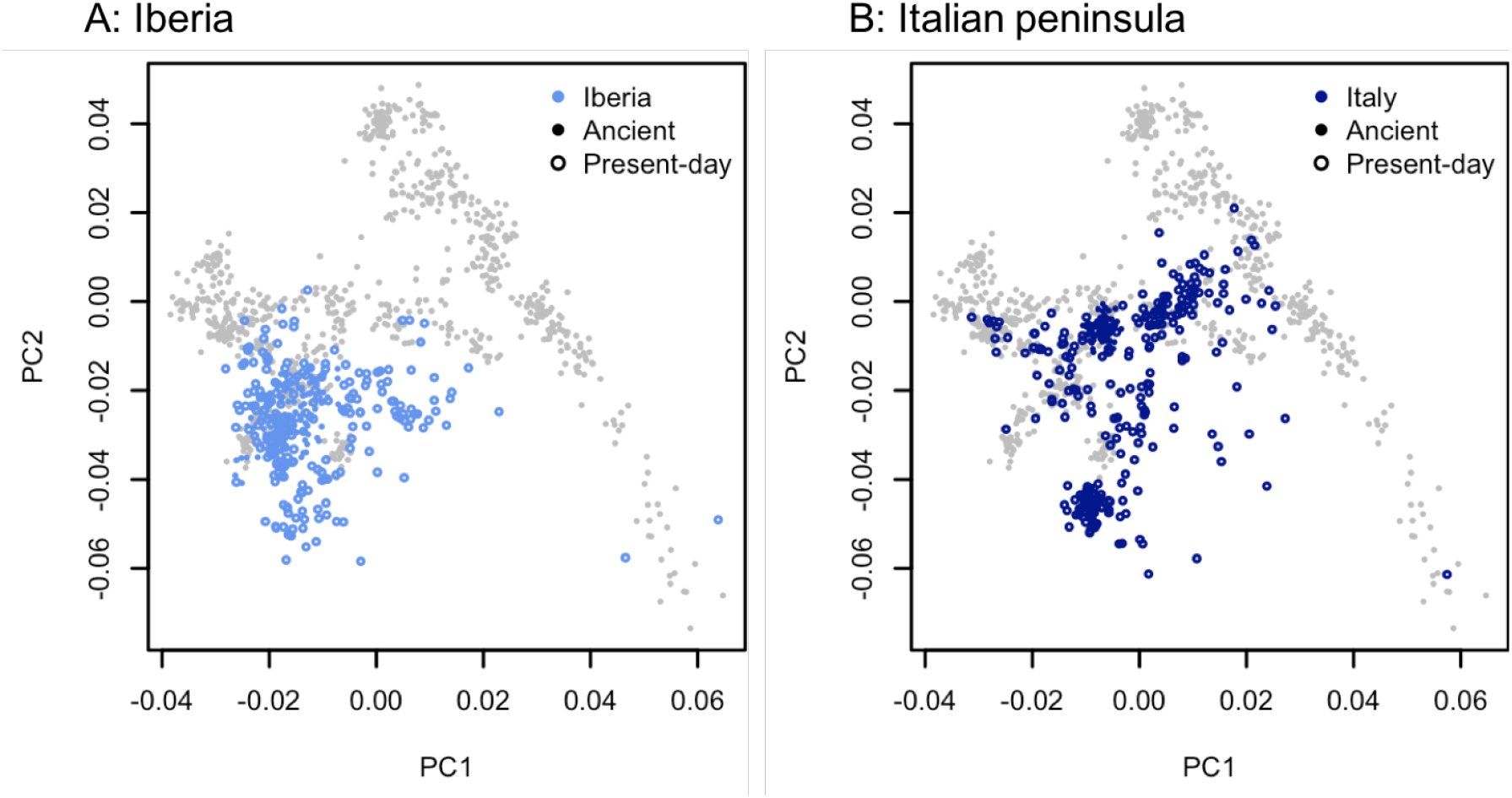
Projected principal components as in Fig. 2 but for ancient individuals from the past 4500 years in Iberia and the Italian peninsula, together with present-day individuals from the IBS and TSI populations of the 1000 Genomes Project. See Ref. [33] for details of the reference populations (grey points).

## Notes

### Competing Interest Statement

The authors have declared no competing interest.

