## Supplemental Notebooks for "Direct detection of natural selection in Bronze Age Britain": bws.py

```
"Beta with spikes (with extensions to selection derived by Paris et al)."

# We encountered various numerical issues with these recursions, such as
# negative values for EZ^2, negative spike probabilities, etc. Running
#
#    $ py.test bws.py
#
# Reproduces some of these issues using the test code included at the end of 
# the file.


import jax
from jax.scipy.special import betaln, gammaln
from jax import numpy as jnp
import numpy as np


def _moment_recursion(mu_t, sigma2_t, s, h, Ne):
    mu_t1 = f_sh(mu_t, s, h) * (1.0 + sigma2_t / 2.0)
    sigma2_t1 = (
        mu_t1
        / Ne
        * (1 - f_sh(mu_t, s, h) - f_sh_prime_prime(mu_t, s, h) / 2.0 * sigma2_t)
    ) + (1 - 1.0 / Ne) * f_sh_prime(mu_t, s, h) ** 2 * sigma2_t ** 2
    return (mu_t1, sigma2_t1)


def _fixation_recursion(
    x0: float, s: float, h: float, Ne: float, t: int
) -> jnp.ndarray:
    "probability of loss or fixation given params"

    def f(carry, _):
        p0, p1, m0, z2_0 = carry
        # unconditional moments
        m, z2 = _moment_recursion(m0, z2_0, s, h, Ne)
        # so E(beta) = m0, var(beta) = z2_0
        # the probability of fixation is (roughly) p(beta) \in [1/Ne, 1-1/Ne] =: E
        # so E(beta') = \int_E x beta(m0, z2_0) / \int_E
        u = 1 - p0 - p1
        m_star = (m - p1) / u
        # z2_star = (z2 + m ** 2 - p1) / (1 - p0 - p1) - m_star ** 2
        z2_star = (
            2 * m * p1 + (-1 + p0) * p1 - m ** 2 * (p0 + p1) + z2 - (p0 + p1) * z2
        ) / u ** 2
        c = m_star * (1 - m_star) / z2_star - 1
        a_star = m_star * c
        b_star = (1 - m_star) * c
        sigma = Ne * s
        p0_prime = p0 + (1 - p0 - p1) * (
            a_star * (1.0 - sigma) + b_star + 2 * Ne
        ) * jnp.exp(
            gammaln(a_star + b_star)
            + gammaln(b_star + 2 * Ne)
            - gammaln(b_star)
            - gammaln(1 + a_star + b_star + 2 * Ne)
        )
        p1_prime = p1 + (1 - p0 - p1) * (
            a_star + b_star + 2 * Ne + b_star * sigma
        ) * jnp.exp(
            gammaln(a_star + b_star)
            + gammaln(a_star + 2 * Ne)
            - gammaln(a_star)
            - gammaln(1 + a_star + b_star + 2 * Ne)
        )
        ret = (p0_prime, p1_prime, m_star, z2_star)
        return ret, None

    init = (0.0, 0.0, x0, 0)
    for _ in range(t):
        init, __ = f(init, _)
    return init
    return jax.lax.scan(f, init, None, length=t)[0]

def f_sh(x: float, s: float, h: float) -> float:
    "allele propagation function for general diploid selection model"
    ret = (
        x
        * (1 + s * h + s * (1 - h) * x)
        / (1 + 2 * s * h * x + s * (1 - 2 * h) * x ** 2)
    )
    # ret = id_print(ret, what="f_sh")
    return ret


def f_sh_prime(x: float, s: float, h: float) -> float:
    return (1 - s * (-2 + x) * x + h * s * (1 + x * (-2 + (2 + s) * x))) / (
        1 + s * x * (-2 * h * (-1 + x) + x)
    ) ** 2


def f_sh_prime_prime(x: float, s: float, h: float) -> float:
    return (
        2
        * s
        * (
            -1
            + x * (3 - s * (-3 + x) * x)
            - 2 * h ** 2 * s * (-1 + x * (3 + x * (-3 + (2 + s) * x)))
            + h * (3 + x * (-6 + s * (3 + x * (-9 + (4 + s) * x))))
        )
    ) / (-1 + s * (2 * h * (-1 + x) - x) * x) ** 3


def _check_fx(*args):
    p0, p1, m, z2 = ans = _fixation_recursion(*args)
    assert z2 >= 0
    assert 0 <= p0 <= 1
    assert 0 <= p1 <= 1
    assert 0 <= m <= 1
    return ans


def test_loss():
    p0, p1, _, _ = _check_fx(0.01, -0.01, 0.5, 1e3, 10)
    assert p0 > 0
    assert p1 == 0.0


def test_fix():
    p0, p1, _, _ = _check_fx(0.99, 0.01, 0.5, 1e3, 10)
    assert p0 == 0.0
    assert p1 > 0.0


def test_extreme():
    p0, p1, _, _ = _check_fx(0.7, 0.1, 0.5, 1e2, 10)
    assert 0 <= p0 < p1 <= 1


def test_drift1():
    p0, p1, _, _ = _check_fx(0.4, 0.0, 0.5, 10.0, 10)
    assert p0 > 0
    assert p1 > 0
    assert p0 > p1


def test_drift2():
    p0, p1, _, _ = _check_fx(0.6, 0.0, 0.5, 10.0, 10)
    assert p0 > 0
    assert p1 > 0
    assert p1 > p0


def test_large_Ne_long_time():
    p0, p1, m, z2 = _check_fx(0.6, 0.1, 0.5, 10000.0, 100)
    np.testing.assert_allclose(p0, 0.0)
    assert p1 > 0


def test_small_Ne_long_time():
    p0, p1, m, z2 = _check_fx(0.6, 0.1, 0.5, 100.0, 100)


def test_symmetry():
    p0a, p1a, _, _ = _check_fx(0.4, 0.0, 0.5, 10.0, 2)
    p0b, p1b, _, _ = _check_fx(0.6, 0.0, 0.5, 10.0, 2)
    np.testing.assert_allclose(p0a, p1b)
    np.testing.assert_allclose(p0b, p1a)


def test_moment_recursion():
    # positive selection, increases mu
    for log_Ne in range(2, 9):
        mu, sigma2 = _moment_recursion(0.5, 0.0, 0.01, 0.5, 10 ** log_Ne)
        assert mu > 0.5
        assert sigma2 > 0.0
    for log_Ne in range(2, 9):
        mu, sigma2 = _moment_recursion(0.5, 0.0, -0.01, 0.5, 10 ** log_Ne)
        assert mu < 0.5
        assert sigma2 > 0.0

    mu = 0.1
    sigma2 = 0.1
    for _ in range(1000):
        mu, sigma2 = _moment_recursion(mu, sigma2, 0.05, 0.5, 1e5)
    assert 1 - mu < 1e-3
    assert sigma2 < 1e-6
    assert sigma2 > 0
    mu = 0.1
    sigma2 = 0.1
    for _ in range(10):
        mu, sigma2 = _moment_recursion(mu, sigma2, -0.05, 0.1, 1e2)
    assert sigma2 > 0


def test_fixation_recursion():
    mu = 0.5
    sigma2 = 0.1
    for _ in range(10):
        mu, sigma2 = _moment_recursion(mu, sigma2, 0.1, 0.5, 1e5)
        assert sigma2 > 0
```
